# Comprehensive public health surveillance identifies novel human cryptosporidiosis subtypes and genetic diversity associated with swimming pool outbreaks in Australia, Victoria (2018-2024)

**DOI:** 10.1101/2025.10.23.684085

**Authors:** Marielle Babineau, Anson V. Koehler, Michelle L. Sait, Karolina Mercoulia, Sally Dougall, Jane McAllister, Evelyn Wong, Norelle L. Sherry, Robin B. Gasser, Benjamin P. Howden

## Abstract

*Cryptosporidium* and *Giardia* are major causes of gastrointestinal illness globally. In Australia, cryptosporidiosis is a nationally notifiable disease, yet molecular characterisation of clinical cases is rarely performed, limiting the capacity to identify outbreaks, trace sources and assess zoonotic risk. During 2024 there was a 273% cases increase in Australia, the third country with the highest increase. We present the first comprehensive molecular investigation of human *Cryptosporidium* and *Giardia* infections in the state of Victoria, Australia. We analysed faecal samples collected between 2018 and 2024. Positive samples were subtyped and parasite load was estimated. Of the 2,330 samples tested, 225 were positive for *Cryptosporidium* and nine for *Giardia*. Seven *Cryptosporidium* species and 24 subtypes were identified, including multiple novel or regionally unique subtypes. *C. hominis* was the predominant species (85%), and three subtypes associated with 11 recreational water outbreaks in 2024. Based on spatiotemporal overlap and subtypes, 52 cases were inferred to represent undetected outbreak-associated infections. Several *C. parvum* subtypes reflected probable zoonotic transmission, two subtypes were associated with a childcare and camp outbreak. Six *C. hominis* subtypes and eight subtypes overall were reported for the first time in Australia. Globally novel subtypes of *C. occultus*, *C. fayeri* and *C. meleagridis* in human hosts is reported. The study reveals high diversity of *Cryptosporidium* subtypes in human infections in Victoria. This study demonstrates the potential for molecular surveillance to inform public health interventions when integrated with epidemiological data. These findings support One Health approaches to outbreak detection and source attribution.

**IMPORTANCE:** *Cryptosporidium* is a nationally notifiable pathogen in Australia, yet routine genotyping is not performed, limiting outbreak detection and source attribution. This study represents the first large-scale molecular surveillance of human *Cryptosporidium* and *Giardia* infections in southern Australia. By integrating species- and subtype-level typing with epidemiological metadata, we demonstrate the substantial diversity of infective lineages, including several novel or zoonotic genotypes. Our findings highlight the critical role of molecular tools in tracking transmission pathways, supporting outbreak investigation and informing public health responses. These data provide a foundation for incorporating routine genotyping into national surveillance strategies for parasitic enteropathogens. This study presents the first epidemiological study of the 2024 outbreaks in Australia. The global comparison of cases during this period also highlights potential large-scale disease dynamic of public health importance.

## INTRODUCTION

*Cryptosporidium* and *Giardia* are among the commonest protistan pathogens causing gastrointestinal illnesses globally, associated with substantial disease burden in both developed and developing countries (1, 2). Cryptosporidiosis is a nationally notifiable disease in Australia, whereas giardiasis is only notifiable in some states and territories, and remains underreported and often unquantified. Recent surveillance studies have documented marked increases in cryptosporidiosis notifications in 2023 and 2024 in Germany, Norway, Belgium, Spain, France and England (3–8). In Australia, an increase in cases linked to swimming pools has been observed, leading to multiple health warnings from public health authorities (9–12). However, the increase of cryptosporidiosis cases in Australia has not yet been comprehensively investigated. A comparison of cases between Australia and other countries during this period has also not been explored. Such a comparison might allow the magnitude of the outbreaks locally and abroad to be quantified, which could identify large-scale patterns of transmission, subtype-specific disease dynamic or common risk factors.

Cryptosporidiosis and giardiasis are primarily transmitted via the faecal–oral route, often through contaminated recreational or drinking water (13, 14). In Australia, infections occur predominantly in children and display marked seasonal peaks during warmer months (15–17). Most reported outbreaks have occurred in urban areas and are linked to recreational water exposure (18–30), although sporadic cases and occasional outbreaks linked to animals (31, 32), drinking water (33–35), unpasteurised milk (36, 37) and overseas travel (38, 39) have also been documented.

Routine clinical diagnosis for *Cryptosporidium* and *Giardia* in Australia relies on qualitative detection in faecal samples using PCR or microscopy, which are unable to differentiate between species or subtypes. There is currently no systematic referral of test-positive samples to reference laboratories for further genetic characterisation, limiting the ability to identify potential outbreaks and to guide targeted public health interventions. This has impeded efforts to resolve transmission pathways, identify outbreak sources and characterise the diversity of human-infective species, genotypes and subtypes. A wide range of *Cryptosporidium* and *Giardia* species and subtypes are known to infect humans, each differing in host range, ecological niche and potential for zoonotic or waterborne transmission (13, 14, 40). To date, 46 *Cryptosporidium* species have been described, with 19 species and more than 60 subtypes reported in humans globally (41, 42). In Victoria, only four species and ten subtypes have previously been recorded from humans (Jex et al., 2007; Koehler et al., 2014). There is no comprehensive study of *Cryptosporidium* subtypes in clinical cases in southern Australia. The identity of *Cryptosporidium* species and subtypes infecting humans remains largely unknown.

Of the eight recognised *Giardia* species, *G. duodenalis* is the principal agent of human infection, with sub-assemblages AI and AII (Assemblage A) and BIII and BIV (Assemblage B) being the most common in humans (40, 45, 46). While these assemblages have been reported in human cases in other states of Australia (47–55), no published data exist on human *Giardia* assemblages in southern Australia, limiting our ability to understand transmission dynamics, identify potential zoonotic sources and assess local public health risk.

In contrast to the limited data for humans, *Cryptosporidium* and *Giardia* species and subtypes have been extensively characterised in non-human hosts across Australia, including wildlife, livestock and companion animals (56–61). In Victoria alone, 24 *Cryptosporidium* species and 31 subtypes have been reported from 16 animal hosts (43, 44, 62–68). *Giardia* assemblages have similarly been recorded in a range of animals in Victoria and nationally (52, 54, 63, 64, 69). This disparity in surveillance limits the assessment of zoonotic risk and obscures the true sources of infection in human populations.

Specific subtypes have been associated with particular transmission routes and risk factors. For example, *C. cuniculus* and *C. fayeri* subtypes have been linked to wildlife contact, and *C. hominis* subtypes IbA10G2, IbA12G3, IgA17 and IdA15G1 are frequently associated with recreational water outbreaks (30, 31, 70, 71), *C. parvum* subtypes are often associated with livestock exposure (71, 72). Accurate species and subtype identification is essential for outbreak resolution, source attribution and the assessment of resistance to antimicrobial drugs. The most widely applied molecular targets for subtyping are the small subunit rRNA (*SSU*) and the 60 kDa glycoprotein (*gp60*) loci for *Cryptosporidium*, and the triosephosphate isomerase (*tpi*) locus for *Giardia* (13, 14, 40). However, these methods have not yet been widely applied to understand outbreaks and transmission, and guide public health policies and interventions.

In this study, we aimed to screen retrospective untested clinical samples to establish the prevalence of these two parasitic infections, we then aimed to subtype all current and retrospective positive clinical samples available, quantify oo/cysts load, and finally integrate subtyping data, parasite load and epidemiology data to identify potential sources and transmission clusters.

## MATERIALS AND METHODS

### Sample set

A total of 2,330 human faecal samples were available to test for *Cryptosporidium* and *Giardia* and genetically characterise these protists (Table 1). A total of 2,329 samples were received at the Microbiological Diagnostic Unit Public Health Laboratory (MDU PHL), Melbourne, between 2020 and 2024, and originated from residents of Victoria (population 6.9 million in 2024 (73)), Australia. Samples were categorised as: (i) *Cryptosporidium*-positive (n = 219), (ii) *Giardia*-positive (n = 9), and (iii) untested samples from symptomatic patients (n = 2,102). One additional *Giardia*-positive sample from 2018 was also included.

**Table 1.**
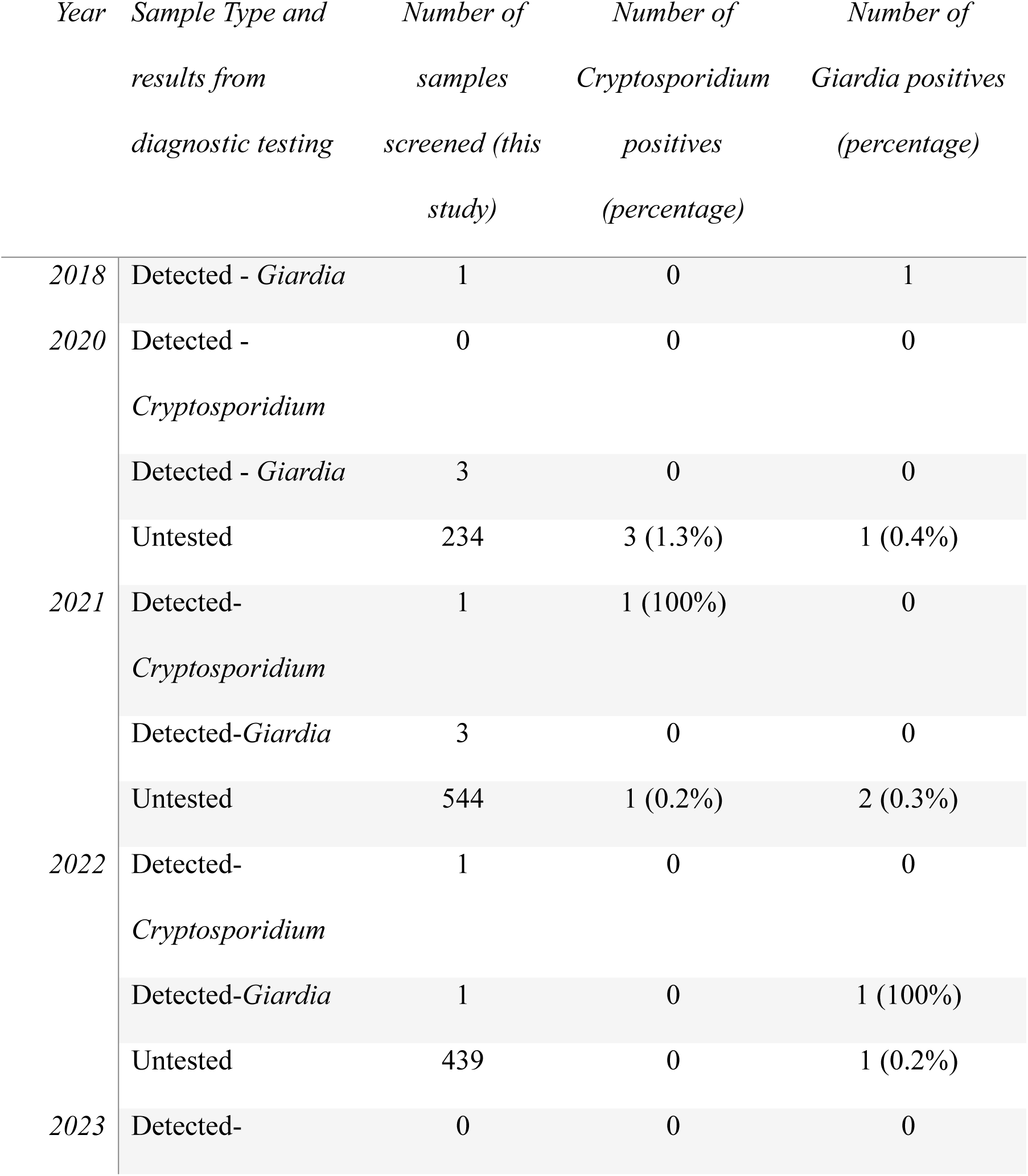
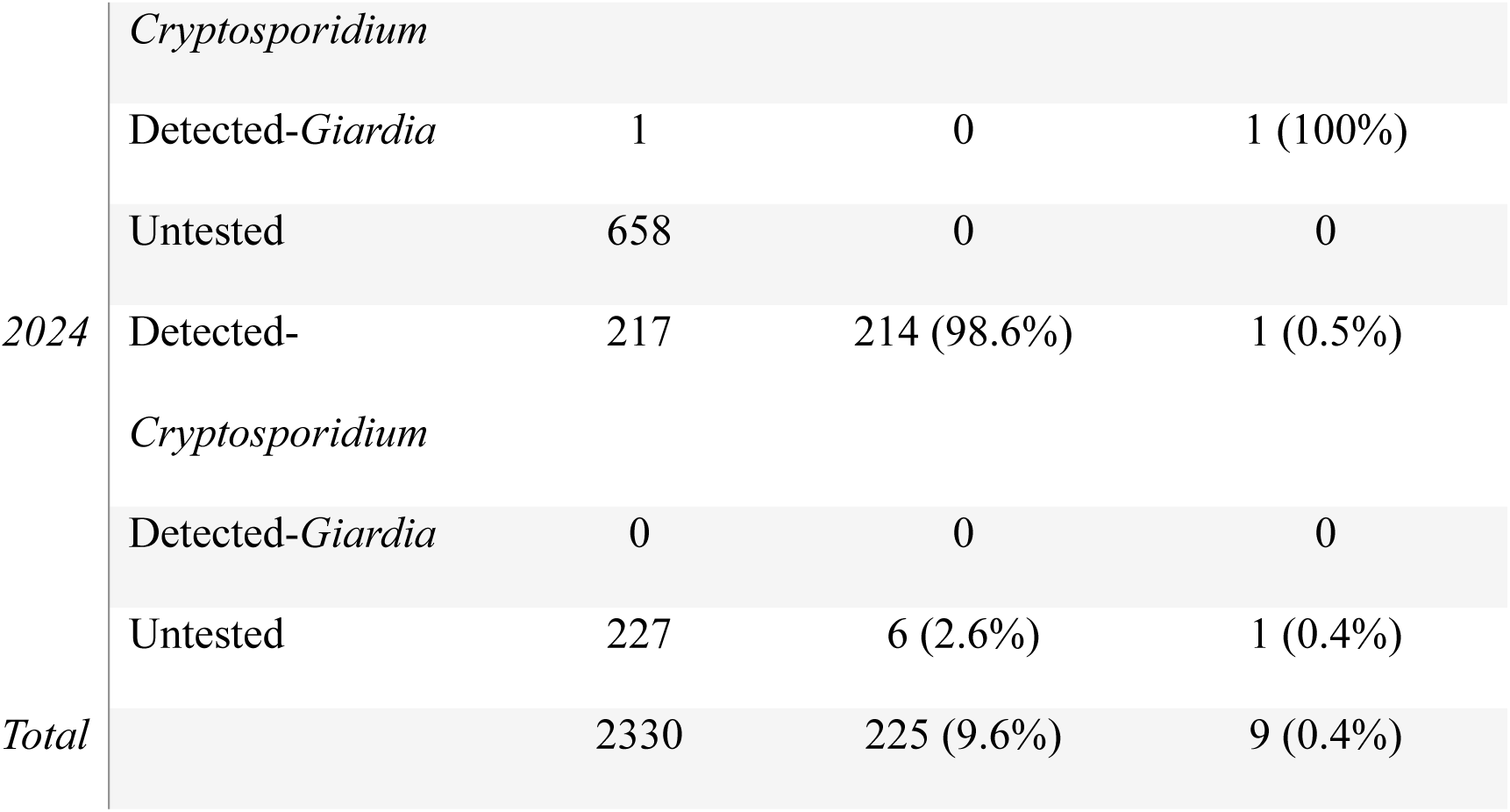
Number of human clinical faecal samples processed by year, sample type, and detection rate of *Cryptosporidium* and *Giardia*. Detected - *Cryptosporidium*; samples in which *Cryptosporidium* was previously detected by PCR or microscopy, Detected - *Giardia*; samples in which *Giardia* was previously detected by PCR or microscopy, Untested; samples from patients with gastroenteric symptoms previously not tested for *Cryptosporidium* or *Giardia*.

*Cryptosporidium*-positive human faecal samples (identified by PCR and/or microscopy) were sent to MDU PHL from multiple diagnostic laboratories. Notified cases are not currently routinely referred for further molecular analysis. Untested samples were submitted from individuals with symptoms of gastroenteritis with the majority of these associated with notified gastroenteritis outbreaks possibly from foodborne or waterborne source, but not previously tested for *Cryptosporidium* or *Giardia*. All samples were stored at –20 °C or –70 °C prior to processing.

### Isolation of genomic DNA

Human faecal samples were thawed at 4 °C and homogenised using a FastPrep®-96 (MP Biomedicals) at 1400 rpm for 10 min. Total genomic DNA was extracted using the QIAamp PowerFecal Pro DNA Kit on the QIAsymphony SP platform (Qiagen, Germany), following the manufacturer’s protocol. Between 0.1–0.4 g of human faecal material (mean: 0.25 g) was used per extraction. To monitor PCR inhibition and DNA recovery, qPCR Extraction Control Red (Meridian Bioscience, USA) was spiked into 1,571 samples across all years. Eluted DNA (85 µl) was stored at – 20 °C. No-template extraction controls were included in each batch (48–96 samples).

### PCR and Sanger sequencing

Nested PCR was used to amplify *Cryptosporidium* small subunit rRNA (*SSU*) and 60 kDa glycoprotein gene (*gp60*) loci, and *Giardia* triosephosphate isomerase (*tpi*) loci, using GoTaq® G2 Hot Start Taq polymerase (Promega, USA) and established protocols (64). *SSU* primers were sourced from previous publications (74–76) and yielded ∼800 bp amplicons. Amplification of *gp60* from *C. meleagridis* or *C. occultus* followed established methods (77, 78). Amplicons were treated with ExoSAP-IT Express (Applied Biosystems, USA) and sequenced bidirectionally using BigDye Terminator v3.1 chemistry (Applied Biosystems). Chromatograms were inspected and trimmed manually using Geneious Prime 2025 v2.1 (79).

### Phylogenetic analysis

Sequences were aligned using MUSCLE (80) in Geneious and refined manually. Sequence identities were assessed via BLASTn using GenBank. Subtypes of *gp60* were assigned according to established nomenclature (78, 81). Substitution models were selected using MEGA v11.0.13 (82) based on the lowest AIC score: GTR+G+I for *SSU* and *gp60*; K2+G+I for *tpi*. Bayesian inference was performed using MrBayes v3.2.7 (83) with 10–50 million generations (depending on locus), four chains, and sampling every 1,000 generations. The first 25% of trees were discarded as burn-in. Convergence was confirmed by split frequency standard deviation (<0.01) and a scale reduction factor of 1.0. Outgroups included *C. muris*, *C. fayeri*, and *G. muris*. Phylogenetic trees were visualised and annotated using iTOL (84).

### Quantification of oocysts and cysts by multiplex qPCR

A validated multiplex qPCR assay (S1. Supplementary Methods) was used to estimate *Cryptosporidium* oocyst and *Giardia* cyst numbers (per gram of faeces) in PCR test-positive samples. The assay targeted *SSU* (*Cryptosporidium*) and the *gdh* gene (*Giardia*) and was calibrated using standard curves from serially diluted DNA derived from *C. parvum* IOWA strain (subtype IIaA15G2R1; Bunchgrass Farms) and *G. duodenalis* (wildtype strain WB1B). Samples were screened in triplicate and samples with Ct values <37 were retained. The mean genome copy number per gram was calculated. The qPCR-based estimates were cross-validated with counts from immunomagnetic separation followed by direct enumeration in a haemocytometer. Assay development and validation protocols are described in the Supplementary Methods.

### Epidemiological data analysis

Epidemiological data were obtained from the Victorian Department of Health for all notified cases of cryptosporidiosis between 2019 and 2024. Data included patient age, sex, hospitalisation status, notification and sample collection dates, outbreak linkage, outbreak setting, and outbreak ID. Age was categorised into 10 years brackets. Cases were categorised as hospitalised if they were admitted to hospital or presented to the hospital emergency. The sample postcodes were matched to local government areas (LGAs) using the Australian Statistical Geography Standard (Australian Bureau of Statistics, July 2011). If a postcode spanned multiple LGAs, the first listed alphabetically was selected. Spatial, temporal and subtype data were used to identify additional cases potentially linked to outbreaks. If a case could be potentially linked to multiple outbreaks, the earliest outbreak was selected. Travel history data and the immune status of the notified cases was not obtained. Statistical correlation between *Cryptosporidium* oocyst load, species, subtypes, and with patient age, sex, hospitalisation status and by outbreak setting were analysed using ANOVA.

### Cryptosporidiosis cases nationally and internationally

The annual summary of cryptosporidiosis cases from 2019 to 2024 was obtained from a total of 32 countries from North America (USA and Canada), Oceania (Australia, with all jurisdictions, and New Zealand), and 28 countries in Europe including the European Union as a whole (S1. Supplementary Methods). Regions with less than ten cases in 2023 or 2024 were excluded. Regions with no data for two or more years in the period were excluded. To the best of our knowledge, the regions identified above are the only ones worldwide where cryptosporidiosis is a reportable disease and have a surveillance system with publicly available data. Regions were classified into three categories based on the increase in cases observed in 2023 or 2024 compared to the previous year; sharp (>50% increase in cases), moderate (15-50% increase), none (< 15%). The percentage of increase for 2023 or 2024 was calculated as: ((latest year case number - previous year case number)/previous year case number)*100. Comparison of the case number for 2023/2024 against the mean of the four previous years was misleading due to global decrease of cases during the COVID-19 pandemic. The comparison of 2023 in the Northern hemisphere with 2024 in the Southern hemisphere reflects the shift in the high-risk summer season across those regions.

### Ethics

Ethical approval to conduct the study was received from the University of Melbourne Human Research Ethics Committee (approval number 2025-30320-66468-5).

### Data availability

Sequences are deposited in NCBI under accession PV981441 - PV981665 (*SSU*), PX092398-PX092617 and PX048014 (*gp60*), PX048005-PX048013 (*tpi*).

## RESULTS

### Detection of parasites in human faecal samples

Of 2,330 human faecal samples tested, 225 were test-positive for *Cryptosporidium* and nine for *Giardia* by PCR-based sequencing (Table 1). One sample from 2024 showed a mixed infection of *Cryptosporidium* and *Giardia*. The detection rate among previously known positive samples was 98%. Detection levels for untested samples ranged from 0% in 2022–2023 to 2.6% in 2024 (Table 1).

### *Cryptosporidium hominis* is the most common species identified among six others

Seven *Cryptosporidium* species were identified using *SSU*: *C. hominis* (n = 191), *C. parvum* (n = 25), *C. meleagridis* (n = 3), *C.* sp. mink genotype (n = 3), *C. fayeri* (n = 1), *C. occultus* (n = 1), and *C.* sp. OTUi (n = 1) (Fig. 1). *SSU* sequences of *C. hominis* showed variation in a T nucleotide repeat region near position 428 bp: 165 had 11T, seven had 8T, and 19 had 9T. The 8T variant also exhibited a downstream Y (C/T) ambiguity at a diagnostic position. Aside from these features, all *C. hominis SSU* sequences were identical. All *C. parvum* sequences were identical to the IOWA strain (OR421305; (85)). One *C. meleagridis* sequence had a single T>A substitution; all three were otherwise identical and matched the UKMEL3 reference (86). The *C.* sp. mink genotype sequences (n = 3, from 2020) were identical and matched two shorter Australian human-derived sequences (87, 88). These human sequences clustered in a polytomy distinct from animal-derived sequences, which grouped with *C. sciurinum*. The conserved nature of the *SSU* locus limited interspecies resolution.

**Figure 1.**
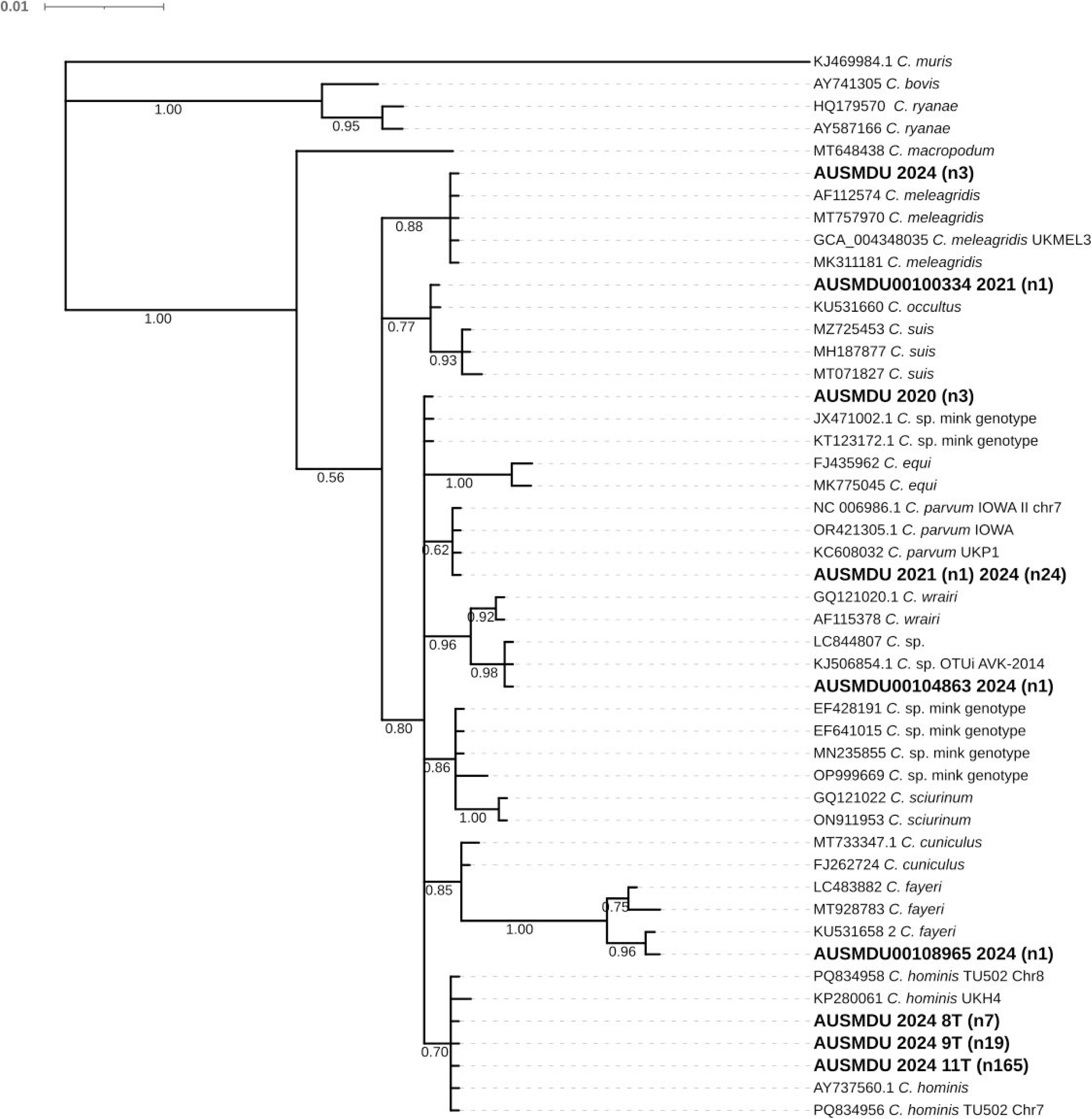
Phylogenetic tree reconstructed from the bayesian topology using GTR +G+I model based on the dereplicated alignment of the small subunit ribosomal RNA gene (*SSU*) with 10M generations for *Cryptosporidium*. Sequences generated in this study are shown in bold with the number of samples identified within parenthesis (n). Posterior probabilities are shown below the branch. *Cryptosporidium muris* used as outgroup.

The *Cryptosporidium* sp. OTUi sequence (2024) matched sequences from a returned traveller and a bat from the Philippines (38, 89) and was 98.9% similar to *C. wrairi*. The *C. fayeri* sequence matched one from a kangaroo in Victoria (64). The *C. occultus* sequence (2021) was identical to that of a deer in Victoria and contained a species-specific motif distinguishing it from *C. suis* (90).

### High intraspecific *gp60* subtype diversity and novelty

Ten *gp60* subtypes were detected among *C. hominis* samples: IaA12R3 (n = 52), IaA14R3 (n = 3), IaA16R3 (n = 1), IaA16R4 (n = 2), IaA24R1 (n = 1), IbA8G4 (n = 1), IdA15G1 (n = 13), IeA11G3T3 (n = 92), IfA12G1R5 (n = 12), and IfA13G1R4 (n = 1) (Fig. 2). Two samples were not amplified. The 9T *SSU* repeat variant was always associated with subtype IeA11G3T3, but this subtype also occurred in 8T and 11T samples. All sequences within each subtype were identical. Several subtypes matched previously reported sequences in Australian or international datasets (e.g., IaA12R3, IdA15G1, IeA11G3T3, IfA12G1R5), while IbA8G4 showed only 90–91% identity to other Australian Ib subtypes. Subtype clades formed strongly supported monophyletic groups by subfamily but lacked resolution at deeper nodes.

**Figure 2.**
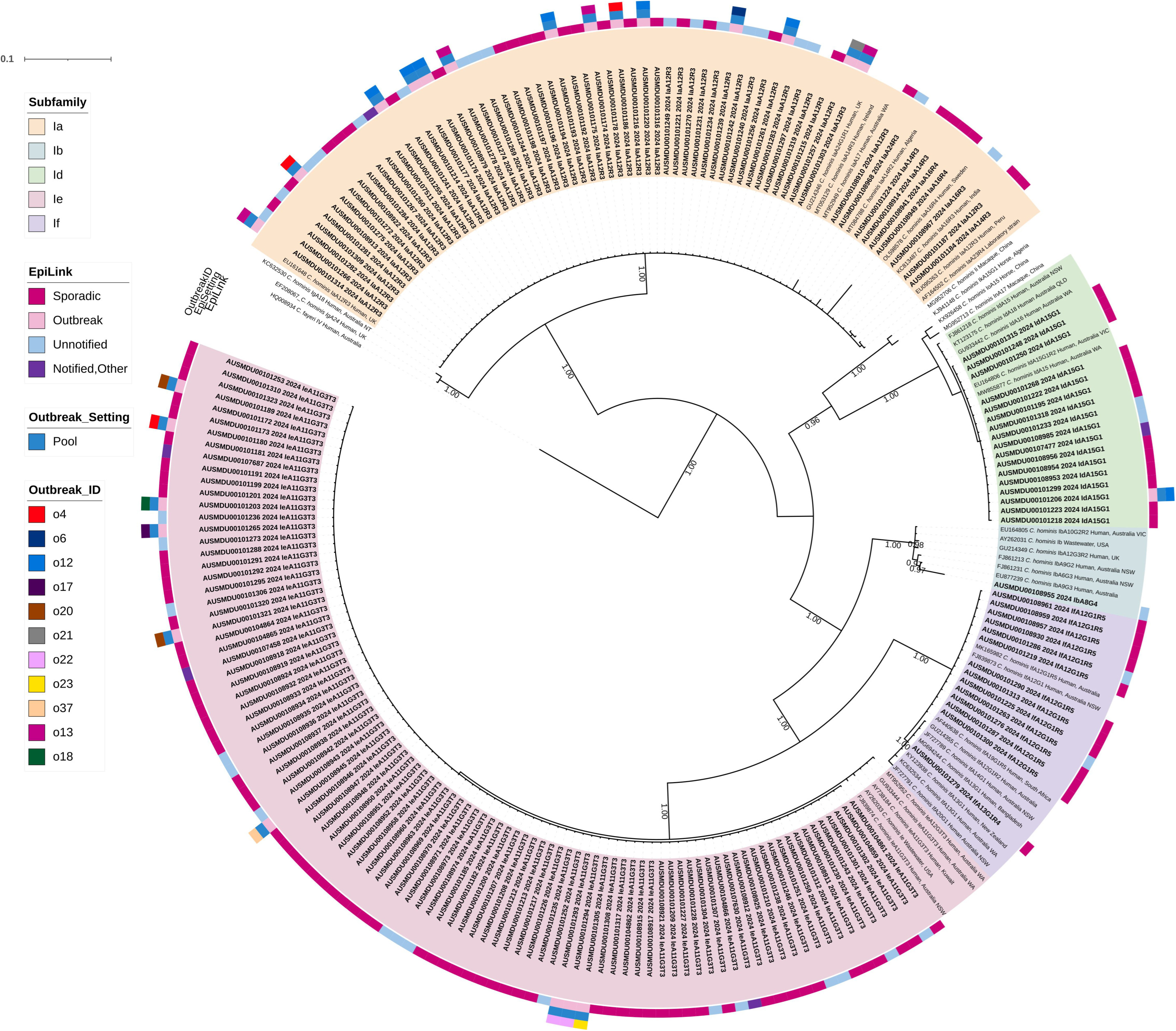
Phylogenetic tree reconstructed from the bayesian topology using GTR +G+I model based on 60kD glycoprotein gene (*gp60*) with 50M generations for *C. hominis* subtypes. Sequences generated in this study are shown in bold. Posterior probability equal to or above 0.90 are shown below the branch. *Cryptosporidium fayeri* used as outgroup. Taxa labels highlighted by *gp60* subfamily identity. Additional data shown as coloured strips; epidemiological link (Epi Link), outbreak setting, outbreak identification number (Outbreak ID). Epi Link: cases classified as sporadic, outbreak related, not notified to Victorian Department of Health as of 18/06/2025.

In the 25 samples containing *C. parvum*, seven *gp60* subtypes were detected: IIaA16G1R1 (n = 1), IIaA16G3R1 (n = 1), IIaA17G2R1 (n = 3), IIaA18G3R1 (n = 9), IIaA19G3R1 (n = 4), IIaA19G4R1 (n = 3), and IIaA20G3R1 (n = 2) (Supplementary Figure S1). Two samples were not amplified. All sequences within each subtype were identical. Most matched cattle- or wildlife-derived Australian sequences (32, 62, 91), while IIaA16G1R1 matched cattle sequences from overseas (92).

Three *gp60* sequences from *C.* sp. mink genotype (subtype A14) were identical to each other and to prior human-derived sequences. One previously published sequence from cattle clustered within the same clade. These sequences were distinct from non-Australian mink sequences, supporting a regional divergence. All non-Australian *C.* sp. mink genotype *gp60* sequences available originate from China. The sequences were more closely related to *C. equi* (Fig. 3).

**Fig 3.**
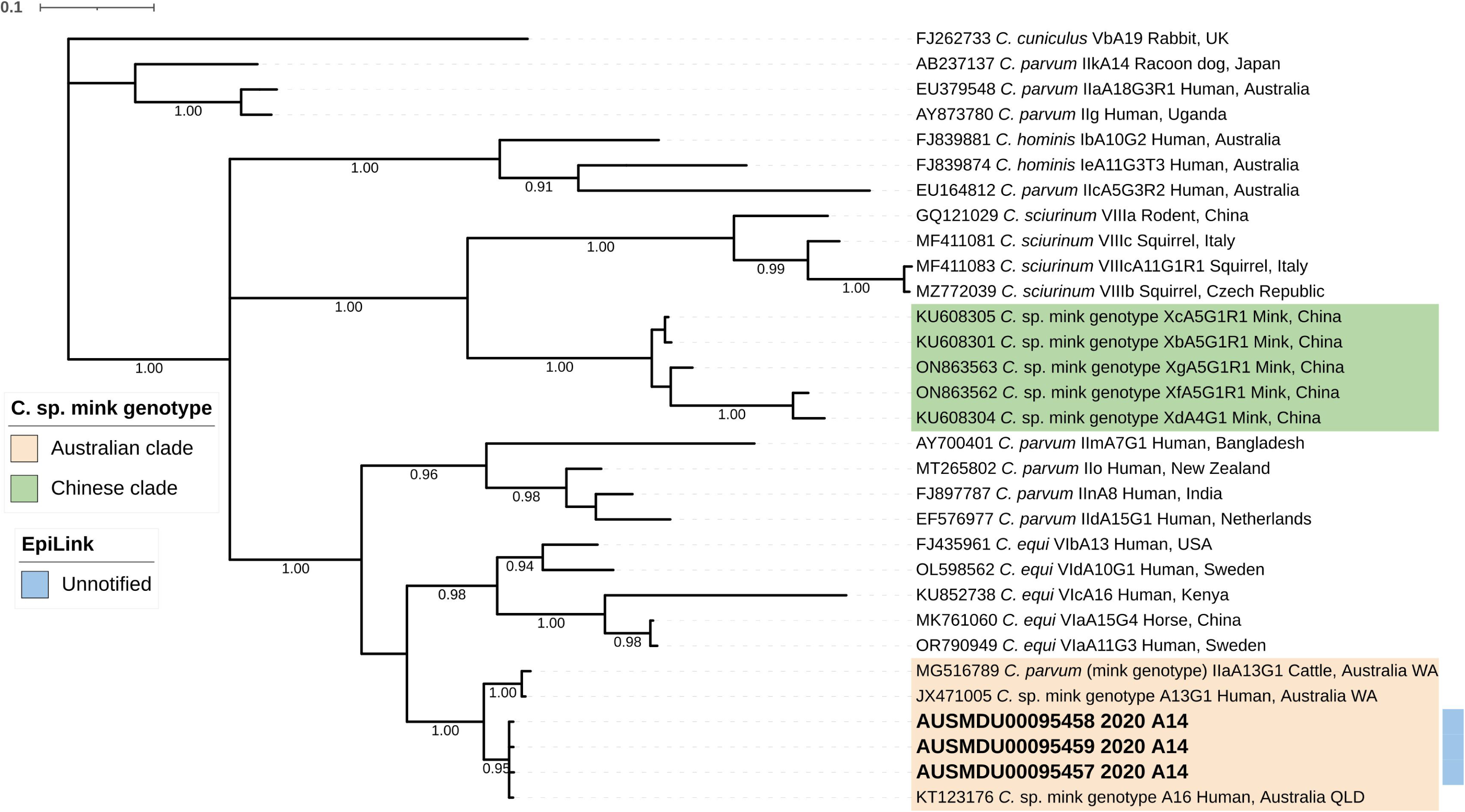
Phylogenetic tree reconstructed from the bayesian topology using GTR +G+I model based on 60kD glycoprotein gene (*gp60*) with 10M generations for *C.* sp. mink genotype sequences. Sequences generated in this study are shown in bold. Posterior probability equal to or above 0.90 are shown below the branch. *Cryptosporidium cuniculus* used as outgroup.

Three *C. meleagridis* subtypes were identified: IIIjA4T1, IIIA19G1, and IIIgA31G4 (Supplementary Figure S2). IIIjA4T1 was 88% similar to a wallaby-derived sequence (93). The IIIA19G1 sequence was not assignable to a known subfamily and contained a unique 33 bp insertion. IIIgA31G4 was 98% similar to a sequence from a human in Sweden (77).

The *C. fayeri* subtype IVfA10G1T1 was 98% similar to sequences from kangaroos (91, 94). The *C.* sp. OTUi *gp60* sequence (subtype A12G1) was 96% similar to a 2014 human-derived sequence (A15G1) and showed a synonymous SNP and a three-repeat deletion. The *C. occultus* subtype was identified as XXIVa and was most similar (94%) to a sequence from a Swedish rat (GenBank accession number PV067140).

### Four *Giardia tpi* assemblages identified

Nine sequences representing *Giardia duodenalis* were assigned to assemblages AI (n = 1), AII (n = 2), BIII (n = 2), and BIV (n = 4) (Supplementary Figure S3). Six were from previously untested symptomatic patients. The AI sequence matched the reference genome strain WB C6 and sequences from a rabbit and kangaroo in Victoria (64). The AII sequences were identical to a previously reported Australian human-derived sequence (88). Four BIV sequences matched samples from Spain and USA wastewater (95). One additional BIV-related sequence clustered within BIII, and two BIII-like sequences (2020 and 2023) showed high similarity (99.1%) to international human-derived sequences.

### Oocyst and cyst load independent of subtypes

Of the 225 *Cryptosporidium* test-positive samples, 194 were subjected to qPCR to estimate parasite load (Table 2). Of the excluded 31 samples, 22 yielded no Ct value and nine had Ct > 37. Oocyst load did not differ significantly by species (F = 0.565, p = 0.758) or subtype (F = 1.247, p = 0.212). Only one *Giardia*-positive sample (from 2018) yielded quantifiable cysts; the remainder failed amplification or exceeded the Ct threshold. Cyst load in the *Giardia* sample was low relative to *Cryptosporidium*. qPCR-based estimates of oocyst load differed by 1–2% from haemocytometer counts following purification via immunomagnetic separation.

**Table 2.**
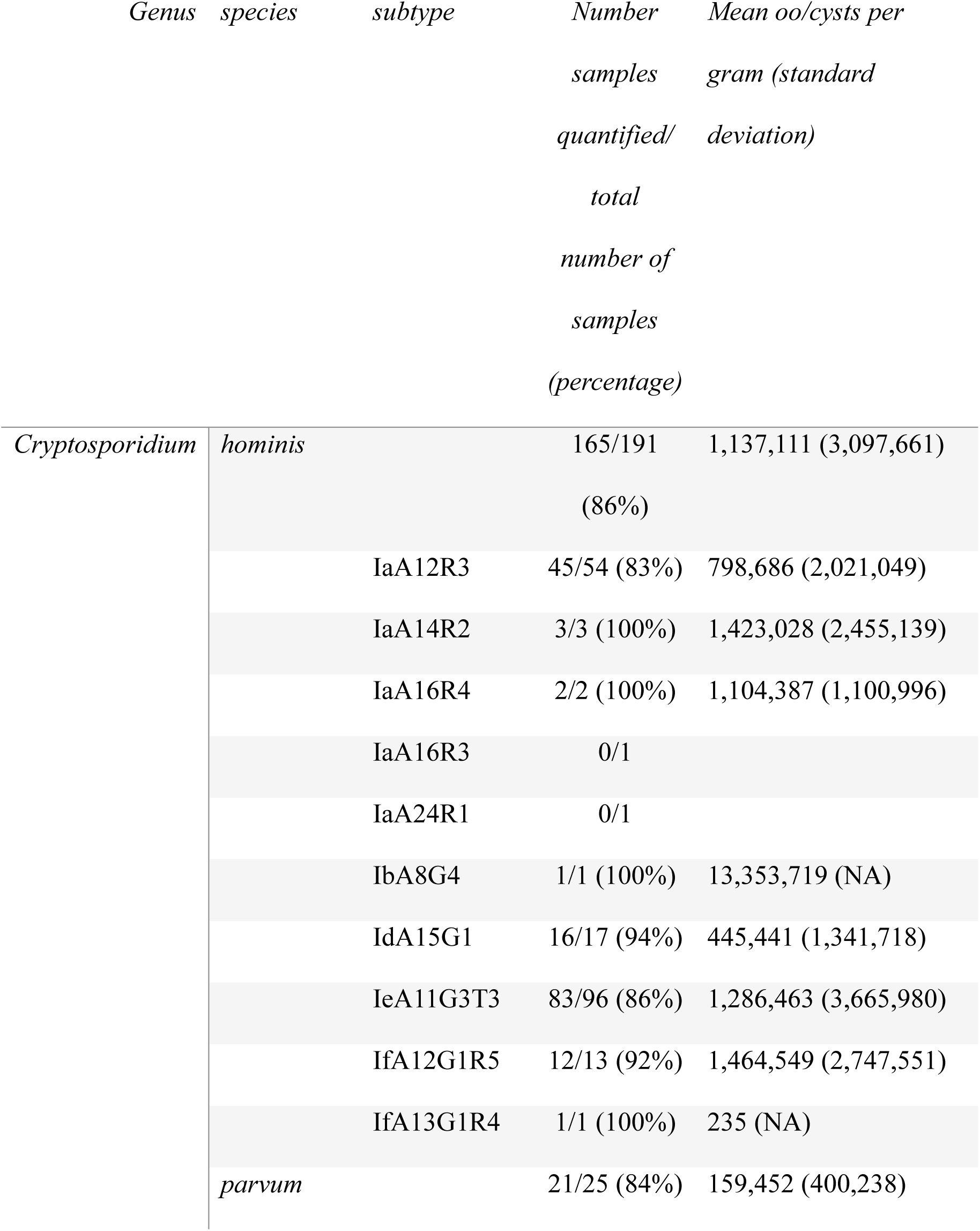
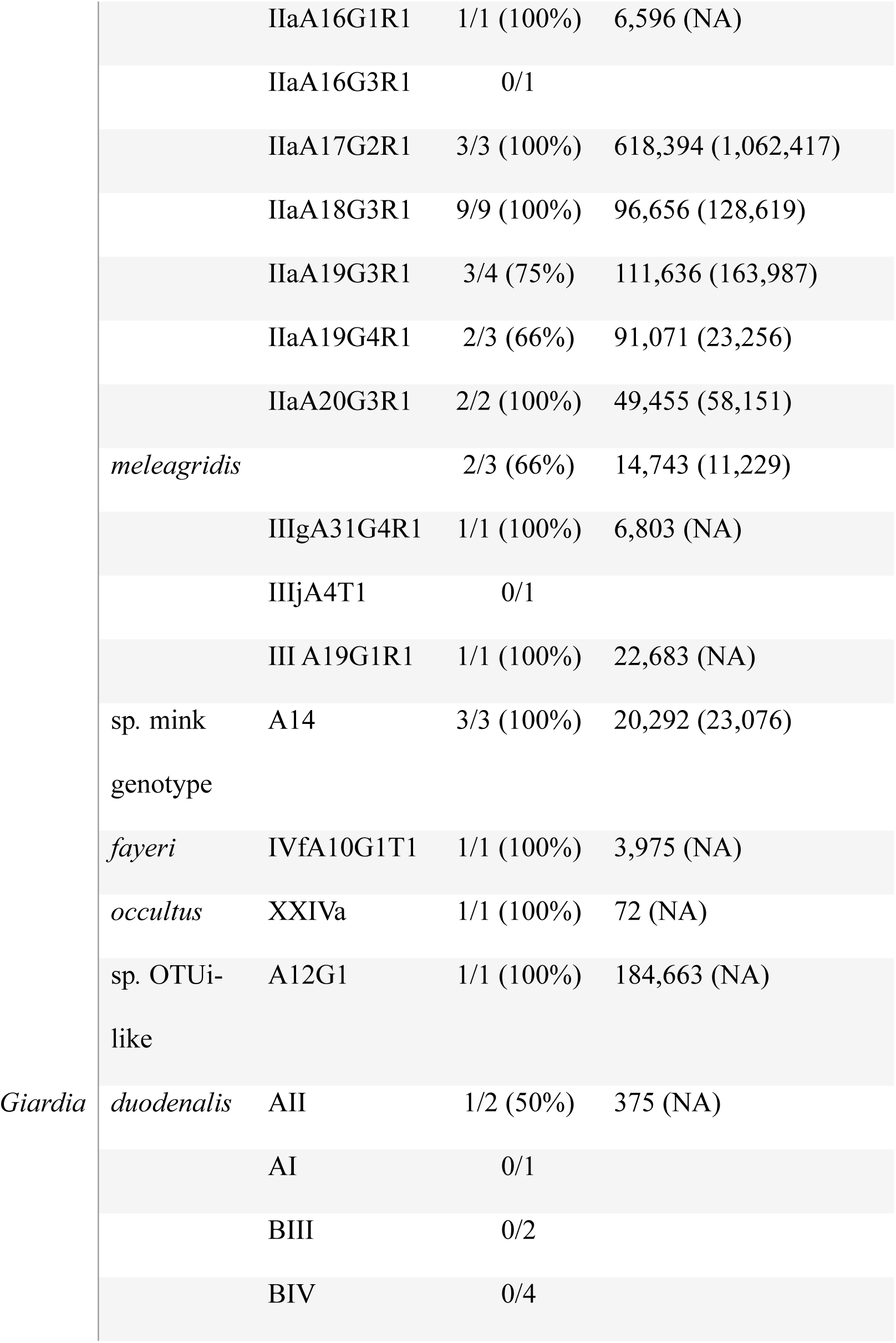
*Cryptosporidium* subtypes and *Giardia* sub assemblages found in this study with respective oocysts and cysts, referred to as oo/cysts, load quantification based on multiplex qPCR. Note: Two *C. hominis* samples and one *C. parvum* sample could not be assigned a subtype but were successfully quantified.

### Subtyping differentiates between swimming pool outbreaks and other outbreak settings

Of 225 *Cryptosporidium*-positive cases, 173 were notified while 52 were previously not notified to public health authorities. Six of the notified cases were notified for a foodborne or waterborne illness other than cryptosporidiosis as they were part of the untested category of samples from 2024, no epidemiological data was available for these six cases. Two cases were positive for rotavirus, one was positive for norovirus while no bacterial or viral pathogens were detected in the other four cases. Four of the non-notified cases, the three *C.* sp. mink genotype and *C. occultus* cases, were newly detected in this study.

The 216 subtyped samples from 2024 represented 6.6% of all notified cryptosporidiosis cases in Victoria in this year (notified between 19 February and 18 September; Fig. 4). Children <10 years accounted for 95 cases, followed by adults aged 31–40 (n = 51). Females comprised 65% (n = 143) of subtyped cases. Subtype identity was not correlated to patient age (F = 0.57, df = 3, p = 0.7).

**Figure 4.**
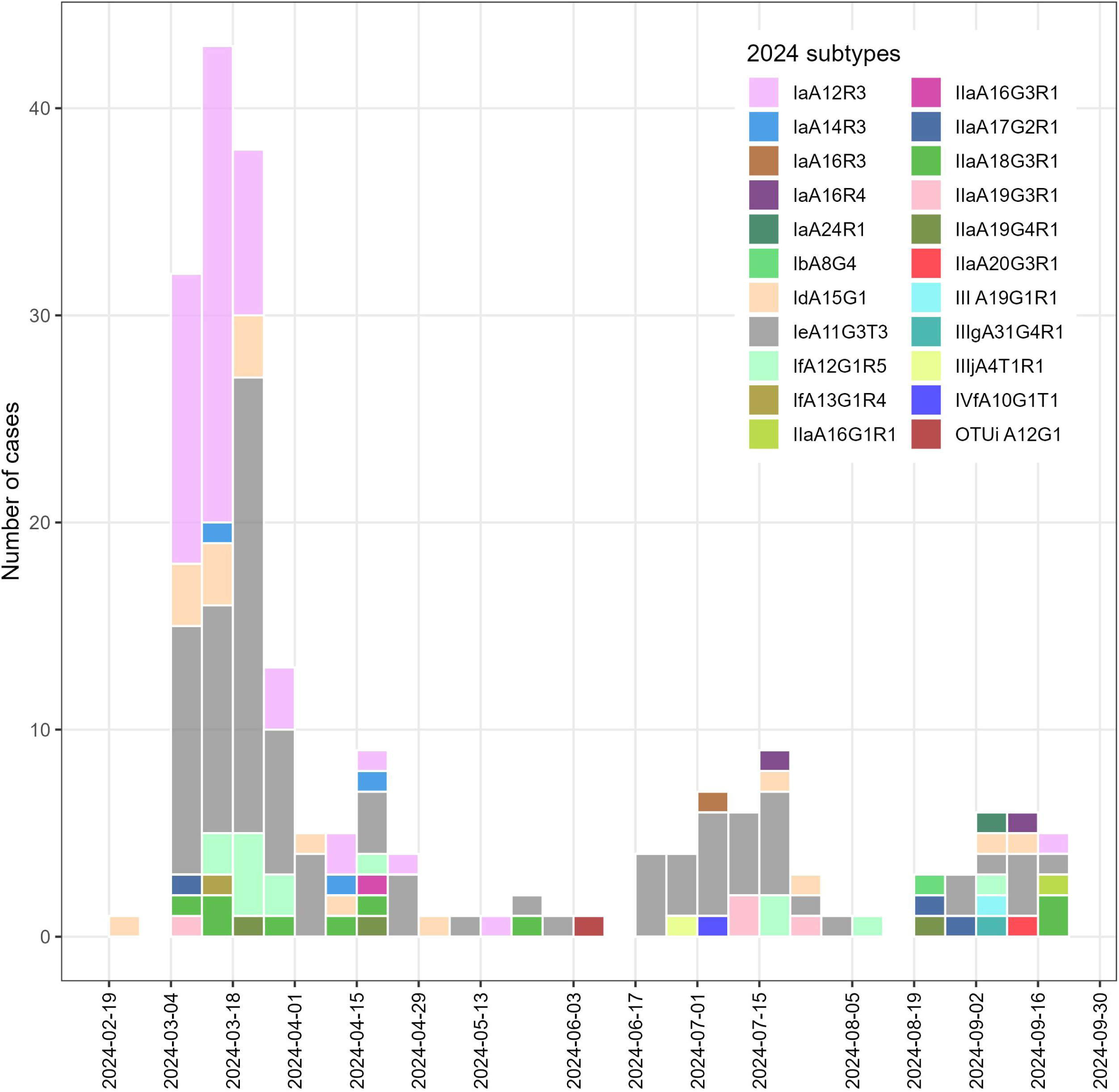
Epidemiological curve of 216 subtyped cases of cryptosporidiosis in 2024 in Victoria.

Among the 167 cryptosporidiosis notified cases with subtype data, 140 (84%) were classified as sporadic and 26 (16%) were epidemiologically linked to 13 outbreaks out of a total of 39 outbreaks ongoing or declared in 2024 in Victoria (11 outbreaks in swimming pools, one outbreak in a childcare, one outbreak in a camp; Fig. 2; Table 3; Supplementary Figure S1). Oocyst load was higher in children <10 years (F = 4.8, df = 201, p = 0.03). Oocysts load was similar between sporadic and outbreak-linked cases, including swimming pool outbreak cases (F = 2.9, df = 2, p = 0.06). No significant difference in oocyst load was observed between sexes (F = 2.1, df = 1, p = 0.15) or hospitalisation status (F = 0.2, df = 4, p = 0.91). Of the 167 cryptosporidiosis notified and subtyped cases, 16 cases (8 females, 8 males) were hospitalized, 15 of which were infected with *C. hominis* - IeA11G3T3 (n=10), IaA12R3 (n=2), IaA14R3 (n=1), IfA12G1R5 (n=1), unknown subtype (n=1) -and the remaining case infected with *C. parvum* (unknown subtype). Only three hospitalized cases were below the age of 10 years old, the majority of subtyped hospitalized cases (n=12) were between the age of 21 to 40.

**Table 3.**
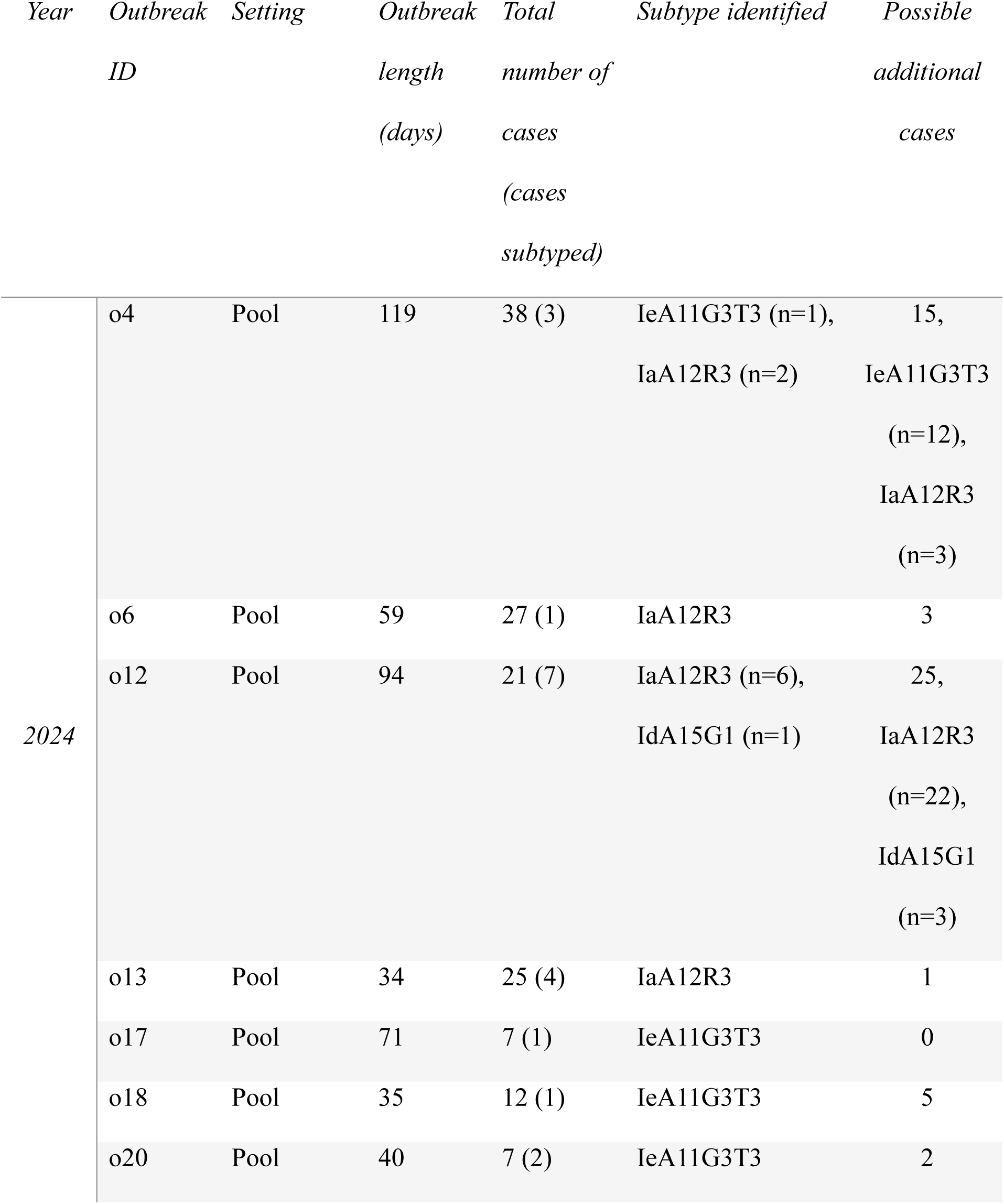
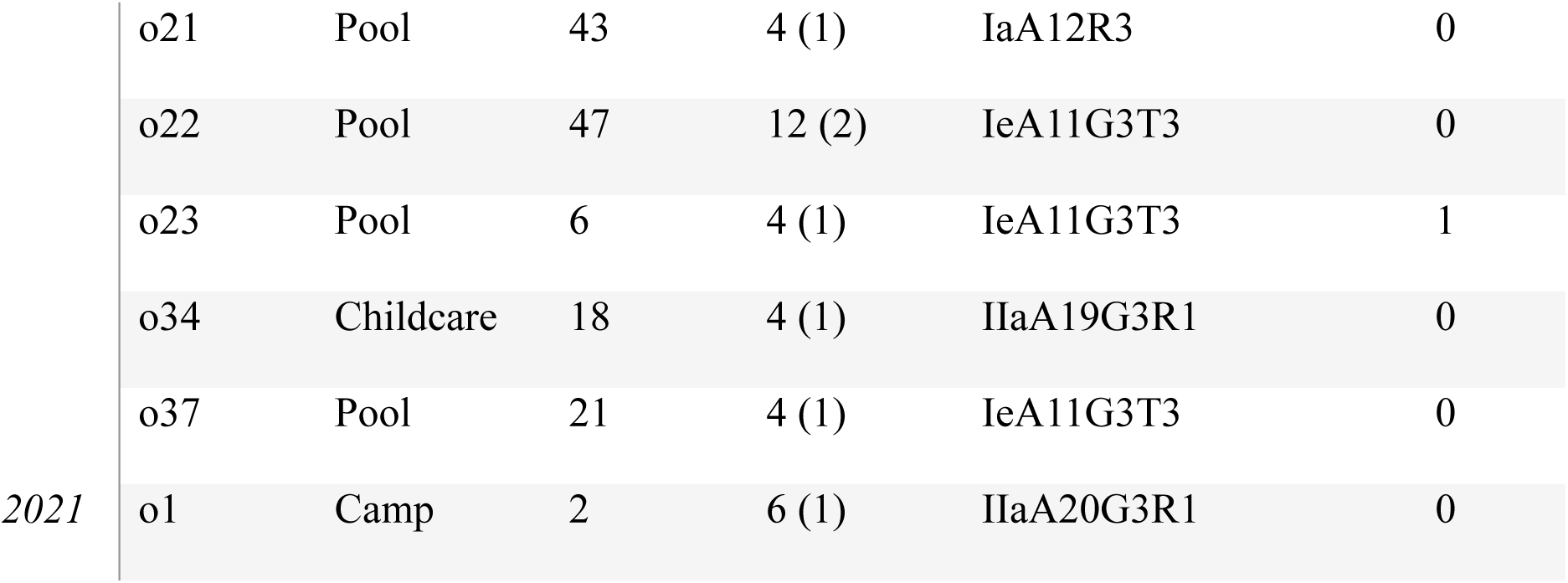
Cryptosporidiosis cases subtyped and epidemiologically linked to outbreaks in Victoria from 2019 to 2024. Possible additional cases are based on cases with the same subtype, notified during the same outbreak period from residents of the same, or directly adjacent, local government areas.

All pool-linked cases were caused by *C. hominis* subtypes IeA11G3T3, IaA12R3 or IdA15G1 (Table 3). The childcare outbreak case involved *C. parvum* subtype IIaA19G3R1 while the case linked to a camp outbreak was *C. parvum* IIaA20G3R1. Two swimming pool outbreaks, o4 and o12, were linked to two subtypes (Table 3). Seven swimming pool outbreaks were linked to subtype IeA11G3T3 and five to IaA12R3. Other subtypes (e.g., IfA12G1R5, IaA16R4, IIIjA4T1R1, IIIgA31G4R1, OTUi A12G1, IIaA18G3R1, IIaA16G1R1) were identified in sporadic cases. Based on spatiotemporal overlap and shared subtypes, 52 additional cases not formally linked to outbreaks were inferred to represent undetected outbreak-associated infections (Fig. 5; Table 3). Two cases could be potentially linked to two different outbreaks; o18 or o20, and another either o4 or o23. The case infected with *C.* sp. OTUi was a female between the age of 21-30. The cases infected by *C. fayeri* and *C. occultus* were not notified and found in a female (61-70 yo) and a male (0-10 yo) respectively, both from the same regional area.

**Figure 5.**
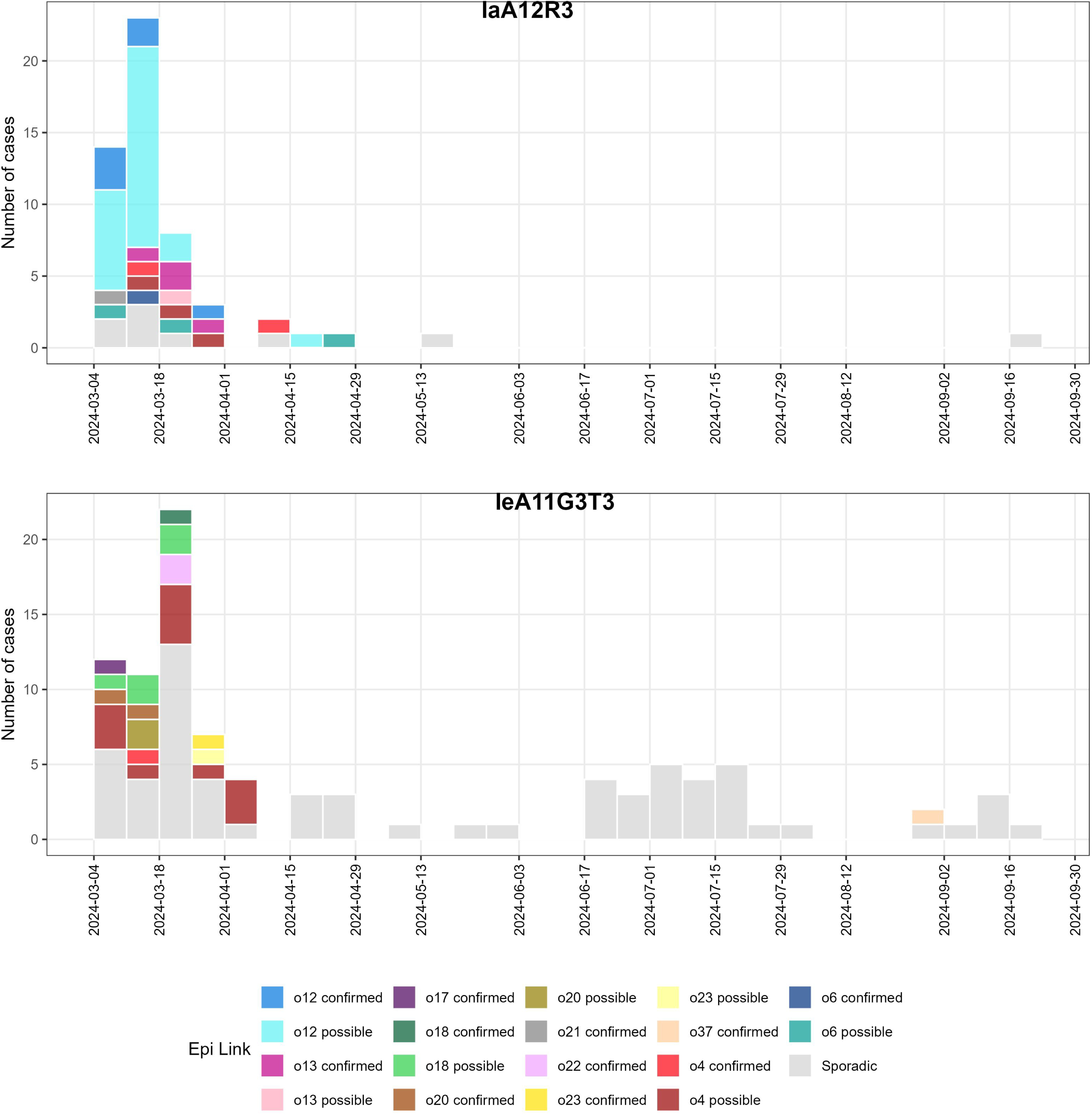
Epidemiological curve of cryptosporidiosis subtype IaA12R3 and IeA11G3T3 cases identified in 2024 in Victoria, classified by epidemiologic links: cases confirmed to be part of an outbreak (e.g. “o12 confirmed” - case part of outbreak o12), cases designated as possibly part of an outbreak, and cases designated as sporadic.

### Australia among the top three countries recording highest rise in cryptosporidiosis cases in 2023-2024

Nationally, Australia recorded a 273% increase in cases in 2024 compared to 2023. Except for the Northern Territory (NT), the number of cases in 2024 from all Australian jurisdictions showed a >200% increase (Fig. 6A). The highest percentage of case increase was in the Australian Capital Territory, Tasmania and Queensland with 793%, 360% and 327% respectively. The highest case numbers were in the Eastern states of Queensland, New South Wales and Victoria. Apart from NT, the number of cases reported in 2024 in all jurisdictions was the highest ever recorded since cryptosporidiosis became notifiable in 2001. Internationally, Australia placed third, behind Iceland and Spain, in countries reporting the sharpest increase in cases in 2023-2024 (Fig. 6B, Supplementary Table 1). A sharp increase in cryptosporidiosis cases in 2023-2024 was also observed in England, Ireland, Belgium, Poland, France, Luxembourg, Slovenia, Malta and Czechia. Australia was the only non-European country to record a sharp increase in cryptosporidiosis during this period. There was no geographical pattern within the European countries which reported a sharp increase in cases. Moderate increases were observed in New Zealand, Canada, Portugal, Latvia, Greece, and Germany. The USA was excluded due to unpublished data for 2023 and 2024.

**Figure 6.**
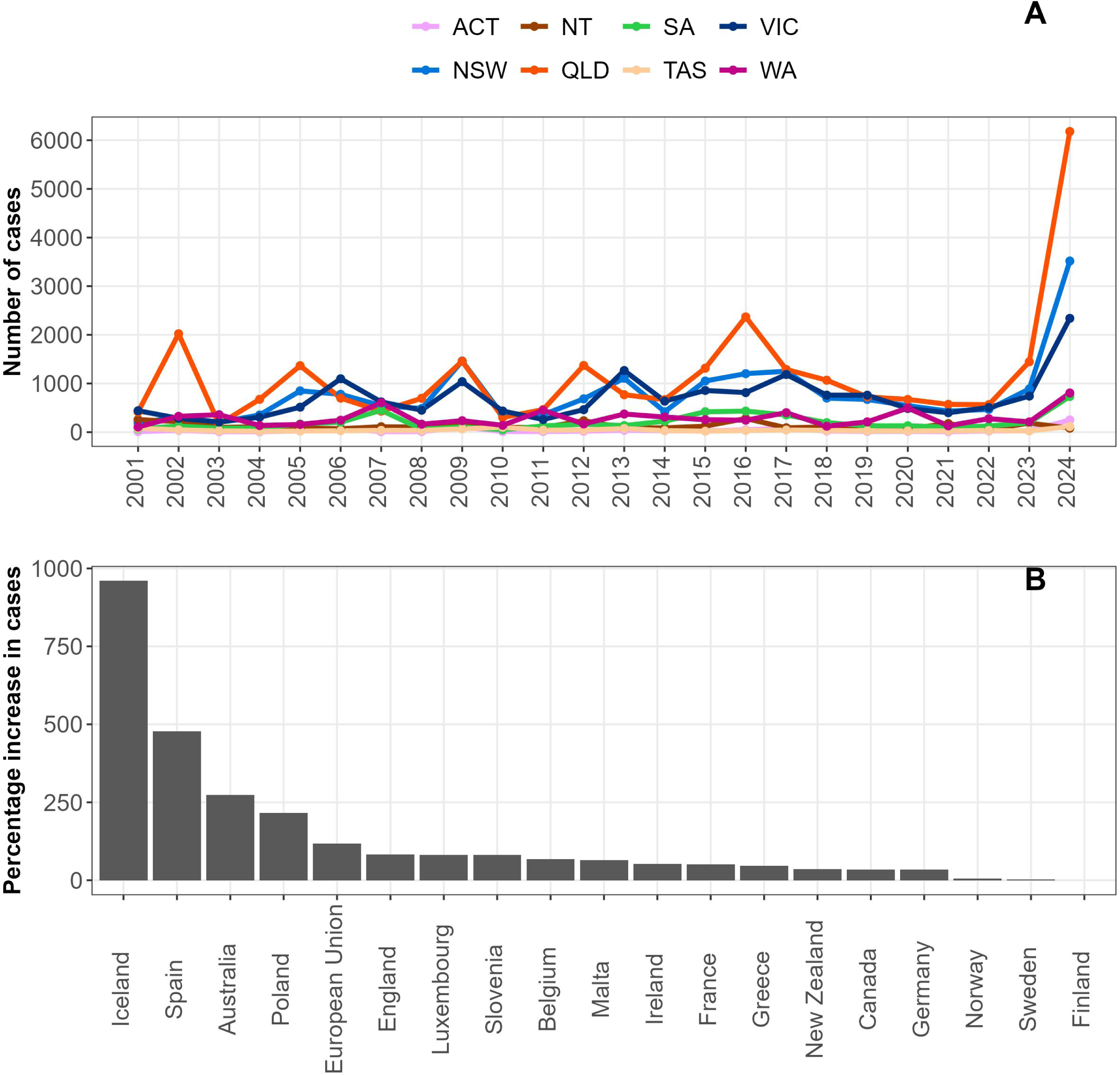
Cryptosporidiosis cases in Australia and abroad. A; annual number of notified cases in each Australian jurisdictions, B; percentage of increase in reported cases from 18 countries (and the European Union as a whole) for the 2023/2024 period compared to the previous year. Abbreviations for Australian states and territories: ACT; Australian Capital Territory, NSW; New South Wales, NT; Northern Territory, QLD; Queensland, SA; South Australia, TAS; Tasmania, VIC; Victoria, WA; Western Australia.

## DISCUSSION

This study presents the first comprehensive molecular investigation of *Cryptosporidium* and *Giardia* infections in people in the state of Victoria, Australia. Of 2,330 faecal samples analysed, 225 were test-positive for *Cryptosporidium* and nine for *Giardia*. The screening of 2,102 untested symptomatic cases did not reveal evidence of large cryptic outbreaks. Low detection rates in 2020–2022 likely reflect reduced transmission under COVID-19 public health restrictions in Australia (96). In 2024, despite multiple cryptosporidiosis outbreaks, there was a very low number of cases, and no evidence of large cryptic outbreaks, related to notified foodborne or waterborne outbreaks based on the untested cases screened in this study.

### Novel taxa within three *Cryptosporidium* species

Three *Cryptosporidium* subtypes identified in this study are taxonomically uncertain and may represent novel lineages. The *C. meleagridis* subtype IIIA19G1 is genetically distinct and not assigned to a known subfamily. The *C. hominis* IbA8G4 sequence differs markedly from the only other published IbA8G4 record and sits on a long branch in the Ib clade, potentially indicating a divergent subfamily. Subtype A12G1 of *C.* sp. OTUi is novel, with a similar subtype from the traveller returning from Indonesia (2014) but no *gp60* sequence from the bat in the Philippines (2025). The *Cryptosporidium* sp. “mink genotype” detected in three patients in 2020 is also distinct compared to other *C. sp*. mink genotype from abroad. The *SSU* and *gp60* sequences from these cases clustered separately from all known animal-derived sequences, supporting the possibility of a regionally divergent Australian lineage. Previous reports in Australians (87, 88) also revealed ambiguous placement in phylogenies. Notably, all three cases here occurred within 48 h during strict COVID-19 lockdown, recorded within different household situated in two suburbs of Melbourne, limiting possible exposure routes. The immune status of these individuals is unknown, one of the cases also tested positive to *Staphylococcus aureus*. Overall, the infection source, transmission route and taxonomic position of this genotype remain unresolved.

### First reports of human infection and subtypes in Australia

This study identified three *Cryptosporidium* subtypes not previously detected in the human host – namely *C. occultus* XXIVa, *C. fayeri* IVfA10G1T1 and *C. meleagridis* IIIjA4T1, previously found only in rats, macropods and rock-wallabies, respectively (78, 93, 94). The identification of *C. meleagridis* IIIjA4T1 in a human raises questions about potential zoonotic or anthroponotic exchange in captive marsupial settings, and the issue of parasite detection in likely passive hosts from zoos and other captive environments. Subtype XXIVa is the first record in a human, previous records being made in animals (mostly rats and a cattle) from Sweden (78).

Additionally, seven subtypes were reported for the first time in Australia, including *C. meleagridis* IIIgA31G4. Six of the ten *C. hominis* subtypes (IaA12R3, IaA14R3, IaA16R3, IaA16R4, IaA24R1, IfA13G1R4) had not been documented in Australia. The highly common subtype IaA12R3 found in this study to be associated with multiple swimming pool outbreaks had not been previously linked to outbreaks worldwide (97). In contrast, all *C. parvum* and *Giardia* subtypes identified herein had been reported previously.

### Subtype dynamics and cryptic outbreak

*Cryptosporidium hominis* was the dominant species (85%), consistent with prior Australian reports (30, 39) and observations from South Asia and sub-Saharan Africa (98, 99). This contrasts with Europe and North America, where *C. parvum* is more common (81, 100–102).

The most frequent subtypes were IeA11G3T3 (41%) and IaA12R3 (23%). While both have been reported previously, neither has been linked to large outbreaks previously (97). Subtype IfA12G1R5, found in 5% of samples in this study, has caused outbreaks in the USA and Spain (5, 103) and was dominant in Western Australia in 2017 (104). Although no outbreak linkage was identified here, its frequency suggests a cryptic transmission cluster. The previously dominant IbA10 subtype was absent, which may reflect evolving strain competition or incomplete sample referral.

Only *C. parvum* subfamily IIa was detected, consistent with zoonotic exposure. Seven subtypes were recorded, several of which (e.g., IIaA16G1R1, IIaA18G3R1, IIaA20G3R1) have been associated with livestock or wildlife in Australia. The absence of subtype IIaA15G2R1, which predominates in Europe and North America (105), further illustrates regional divergence in subtype distribution.

### Zoonotic and environmental infection risk

Multiple subtypes identified here have previously been reported in a restricted host range in Australia (e.g., *C. occultus* in deer, IVfA10G1T1 and IIaA19G4R1 in kangaroos, IIaA16G1R1 in cattle). Internationally, these same subtypes have been detected in diverse hosts, suggesting broad zoonotic potential (41, 106). Thus, local infections may arise via zoonotic or environmental transmission or overseas acquisition. Ongoing One Health surveillance across humans, animals and environments is essential to clarify source attribution.

Molecular surveillance should incorporate spatiotemporal and epidemiological data, including travel history. The absence of travel information in this dataset limits source inference. The case of *C.* sp. OTUi, previously reported in a traveller from Indonesia and a Southeast Asian bat, highlights the importance of contextualising findings with travel and wildlife exposure.

### Quantification and transmission dynamics

Oocyst loads estimated by qPCR were consistent with previous reports from Australian patients (107, 108). No species or subtype-specific differences in load were observed, suggesting that all are capable of high shedding and transmission. Higher loads were observed in children, supporting prior studies indicating increased susceptibility and transmission potential in young age groups (109, 110). Oocysts loads did not appear to be a marker of disease severity based on hospitalisation data available. However, hospitalisation is likely to be under-reported as not all sporadic cases could be followed up due to limited resources at the time.

### Outbreaks and public health risk

Recurring outbreaks linked to swimming pools are well documented in Australia (18–30). The continued detection of *C. hominis* subtypes in pool-linked outbreaks here highlights persistent issues in infection prevention and water quality control. Risk factors include pool design, filtration efficacy, hygiene compliance and microbiological monitoring (111, 112).

Subtype overlap across geographically dispersed outbreaks suggests wider or sustained transmission. Notably, IeA11G3T3 and IaA12R3 were involved in 11 outbreaks and also identified in many other sporadic cases from adjacent LGAs during the same period, suggesting potential under-detection of outbreak scale. The detection of two different subtypes within outbreaks suggests either cases belonging to two different outbreaks or a mixed population as outbreak initial infection source. The two outbreaks linked to *C. parvum* suggest a primary zoonotic infection source and possibly secondary anthroponotic transmission between children. Subtype IIaA19G3R1 and IIaA20G3R1 have previously been recorded in Victoria from deer, alpaca and cattle hosts (62, 113, 114). Many *C. parvum* subtypes (e.g., IIaA18G3R1, IIaA19G4R1) occurred in multiple sporadic cases and may represent cryptic zoonotic outbreaks. Strengthening the One Health approach would improve identification of such linkages.

This study presents the first Australian molecular and epidemiological data from the 2023- 2024 period which saw multiple outbreaks across Australia and the highest number of cases recorded in the last 24 years. Additional data from that period from other jurisdictions would allow further assessment of the scale of *Cryptosporidium* transmission and identify common sources of infection. Previous reviews have found a greater number of *Cryptosporidium* outbreaks from North America and New Zealand (115, 116) compared to other regions suggesting a different disease dynamic during the 2023/2024 period, however the lack of data from the USA limits interpretation from North America. The role of swimming pools in other Australian jurisdictions has yet to be quantified and compared to the Victorian data. The sharp increase in cases observed in several European countries suggest common risk factors specifically increased during this period. The European studies published indicated outbreaks also linked to swimming pools and international travel (4–8). The rise of cases globally during the 2023-2024 summers could be due to this period being one of the warmest on record (117–119). Cryptosporidiosis infections have been shown to increase in warmer months (16, 101, 120) suggesting an increasing risk from this disease in the future. The association between increased temperature and increased cryptosporidiosis cases linked to swimming pools indicate a need for public health authorities to increase prevention and surveillance of cryptosporidiosis in this setting globally. The association with international travel and the worldwide increase in cases suggests that outbreaks could have an increasingly international scale. Worldwide transmission of *C. parvum* subtype has been documented previously (105).

### Genotyping methods and future directions

The *gp60* gene is the most widely used subtyping marker but has limited discriminatory power for outbreak resolution (103, 121, 122). Higher-resolution tools, including MLVA (123), multilocus sequence typing (124), RNA-bait (125), and whole genome sequencing (105, 121, 126), are increasingly being used. However, few are routinely applied to the full taxonomic range of *Cryptosporidium* infecting humans. Whole genome sequencing offers the highest resolution but is not yet adopted for routine typing. Given its status as a nationally reportable disease in Australia, PCR-based subtyping tools, employing *SSU* and *gp60* as markers, are already available, practical and should be systematically applied to notified cases of cryptosporidiosis.

## Conclusion

This study identified seven *Cryptosporidium* species and 24 subtypes, and four *Giardia* assemblages from human clinical samples in Victoria over a five-year period. The observed diversity and subtype distribution suggest multiple transmission pathways – including undetected outbreaks and zoonotic reservoirs – operating concurrently. High sequence identity with animal and environmental isolates supports the need for integrated One Health surveillance. The subtyping of clinical samples, linked to metadata, provides a powerful foundation for improving cryptosporidiosis and giardiasis control and prevention in Australia.

## SUPPLEMENTAL MATERIAL

Supplementary Methods

Supplementary Table 1

Supplementary Figure S1, S2, and S3

## ACKNOWLEDGEMENTS

We thank Louise Baker (Walter and Eliza Hall Institute of Medical Research) for providing the purified *Cryptosporidium* oocysts and *Giardia* trophozoites. We thank all the Victorian microbiology laboratories who forwarded positive *Cryptosporidium* samples in 2024, and the laboratory technical staff at Microbiological Diagnostic Unit Public Health Laboratory for sample reception and pre-processing. We thank Damien Costa and Loic Favennec (University Hospital of Rouen) and the French National Network on Surveillance of Human Cryptosporidiosis for providing case data for France. We thank Guy Robinson (*Cryptosporidium* Reference Unit, Public Health Wales Microbiology) for providing subtyping technical information in relation to *C. occultus*. Funding provided by the University of Melbourne’s co-contribution to Australian Pathogen Genomics Program, funded by the Medical Research Future Fund Genomics Health Future Mission – Pathogen Genomics Grant FSPGN00049 (B.P.H.) and from R.B.G.’s research program which include the Australian Research Council (ARC; LP220200614 and LP180101085) and Oz Omics Pty Ltd.

